# A spiking neural program for sensory-motor control during foraging in flying insects

**DOI:** 10.1101/2020.08.10.243881

**Authors:** Hannes Rapp, Martin Paul Nawrot

## Abstract

Foraging is a vital behavioral task for living organisms. Behavioral strategies and abstract mathematical models thereof have been described in detail for various species. To explore the link between underlying neural circuits and computational principles we present how a biologically detailed neural circuit model of the insect mushroom body implements sensory processing, learning and motor control. We focus on cast & surge strategies employed by flying insects when foraging within turbulent odor plumes. Using a spike-based plasticity rule the model rapidly learns to associate individual olfactory sensory cues paired with food in a classical conditioning paradigm. We show that, without retraining, the system dynamically recalls memories to detect relevant cues in complex sensory scenes. Accumulation of this sensory evidence on short time scales generates cast & surge motor commands. Our generic systems approach predicts that population sparseness facilitates learning, while temporal sparseness is required for dynamic memory recall and precise behavioral control. Our work successfully combines biological computational principles with spike-based machine learning. It shows how knowledge transfer from static to arbitrary complex dynamic conditions can be achieved by foraging insects and may serve as inspiration for agent-based machine learning.

---

Navigating towards a food source during foraging requires dynamical sensory processing, accumulation of sensory evidence and appropriate high level motor control. Navigation based on an animals olfactory sense is a challenging task due to the complex spatiotemporal landscape of odor molecules. A core aspect of foraging is the acquisition of sensory cue samples in the natural environment where odor concentrations vary rapidly and steeply across space. Experimental access to the neural substrate is challenging in freely behaving insects. Biologically realistic models thus play a key role in investigating the relevant computational mechanisms. Consequently, recent efforts at understanding foraging behavior have focused on identifying viable computational strategies for making navigational decisions (1).

An odor plume is often considered a volume wherein odor concentration is generally above some behavioral threshold. At macroscopic scales and in a natural environment, however, plumes are turbulent (2, 3). In turbulent conditions a plume breaks up into complex and intermittent filamentous structures that are interspersed with clean air pockets or below behavioral threshold concentration patches (4, 5). The dispersing filaments form the cone-like shape of the macroscopic plume where the origin of the cone yields the position of the odor source. When entering the cone, flying insects encounter odor filaments as discrete, short-lived sensory events in time.

Several features have been derived from the statistics of an odor plume that provide information regarding the location of the odor source (3, 4). The mean concentration varies smoothly in lateral and longitudinal directions of time-averaged (and laminar) plumes. However, for behavioral strategies animals cannot afford the time it takes to obtain stable macroscopic estimates of mean concentrations (2). Hildebrand and colleagues (6) proposed the time interval between odor encounters as an informative olfactory feature while (3) suggested intermittency, the probability of the odor concentration being above some behavioral threshold, as the relevant feature. However, similarly to estimating mean concentration, acquiring a sufficient number of samples for stable estimates of these quantities exceeds the time typically used to form behavioral decisions (2). Hence, obtaining time averaged quantities is not an optimal strategy to guide navigational decisions as concluded by (7).

Most animals perform searches at large distances from the odor source where the intermittency of plumes poses a more severe problem as available sensory cues become more sparse in space and time. Thus, strategies that exploit brief, localized sensory cues for navigation have been studied by several groups. One strategy for medium and long-range navigation that has consistently been observed across species of flying insects emerges from a sequence of chained sensori-motor reflexes: casting & surging (8). Encountering a whiff of odor triggers an upwind surge behavior, during which the insect travels parallel to the wind direction. After losing track of the plume it evokes a crosswind cast behavior, in which a flight path perpendicular to the direction of air flow is executed. Performing repeated casts by U-turning allows the insect to reenter and locate the plume in order to trigger the next upwind surge (8–10). As the subject approaches the source it increasingly makes use of visual cues for navigation as the plume narrows down. (8).

A number of studies have proposed abstract mathematical models for optimal search algorithms that assumed different types of relevant navigational cues. The infotaxis method proposed in (11) depends on extensive memory and priors regarding a plume’s structure. Contrary, in (8) only local cues are used. A standard algorithm for navigational problems in robotics is simultaneous localisation and mapping (SLAM), which has been used in (12) to study olfactory navigation in bumblebees. An algorithm that works without space perception has been proposed by (13) using a standardized projection of the probability of source position and minimization of a free energy along the trajectory. Finally, the work of (10) compares several models and shows that it is difficult to discriminate between different models based on behavioral responses. A recent work by (7) using information-theoretic analysis shows that plumes contain both, spatial and temporal information about the source’s position.

While all of these previous mathematical methods for olfactory search algorithms have proven to successfully solve this task based on the respective assumptions, they share the same major drawback: none of them uses the computational substrate of the brain, spiking neurons and networks thereof. Instead, all methods make heavy use of symbolic math and advanced mathematical concepts that are not available to the biological brain. It is further unclear how and to what extend these methods could be implemented or learned by the nervous system. Additionally, the problem of navigation and foraging is often considered as an isolated task, independent from sensory processing.

Our approach distills recent experimental results to formulate a biologically plausible and detailed spiking neural network model supporting adaptive foraging behavior. We thereby take advantage of the rapidly accumulating knowledge regarding the anatomy (e.g. (14–16)) and neurophysiology (e.g. (17–19)) of insect olfaction and basic computational features (20, 21). We follow the idea of compositionality, a widely used concept in mathematics, semantics and linguistics. According to this principle, the meaning of a complex expression is a function of the meanings of its constituent expressions (Frege principle (22)). In the present context of foraging and navigation this means dynamically recombining memories of individual sensory cues present within a plume.

## Results

We approach the problem of foraging by decomposition into four components: First, sensory processing with temporal sparse and population sparse coding in the mushroom body (MB). Second, associative learning for assigning a valence to individual odor identities. Third, the time-dependent detection of valenced cues resulting from encounters of discrete odor filaments to provide an ongoing and robust estimate of sensory cue evidence. Fourth, the translation into online motor command signals to drive appropriate behavior.

For sensory processing we use a three-layer spiking neural network model of the insect olfactory pathway (see Fig 1). The generic blueprint of the insect olfactory system is homologous across species and comprises three successive processing stages (see Materials and Methods for details): The periphery with olfactory receptor neurons (ORNs), the antennal lobe (AL) and the MB. Excitatory feed-forward connections across layers from ORNs to projection neurons (PNs), from ORNs to local interneuron (LNs), and from PNs to the MB Kenyon cells (KCs) are fixed. Lateral inhibition within the AL uses fixed synaptic weights from LNs to PNs. For neuron numbers and their connectivity patterns we here rely on the adult *Drosophila melanogaster* where anatomical knowledge is most complete (14, 23, 24). A single MB output neuron (MBON) receives input from all Kenyon cells and plasticity at the synapses between KCs and the MBON enables associative learning (25, 26).

**Fig. 1.**
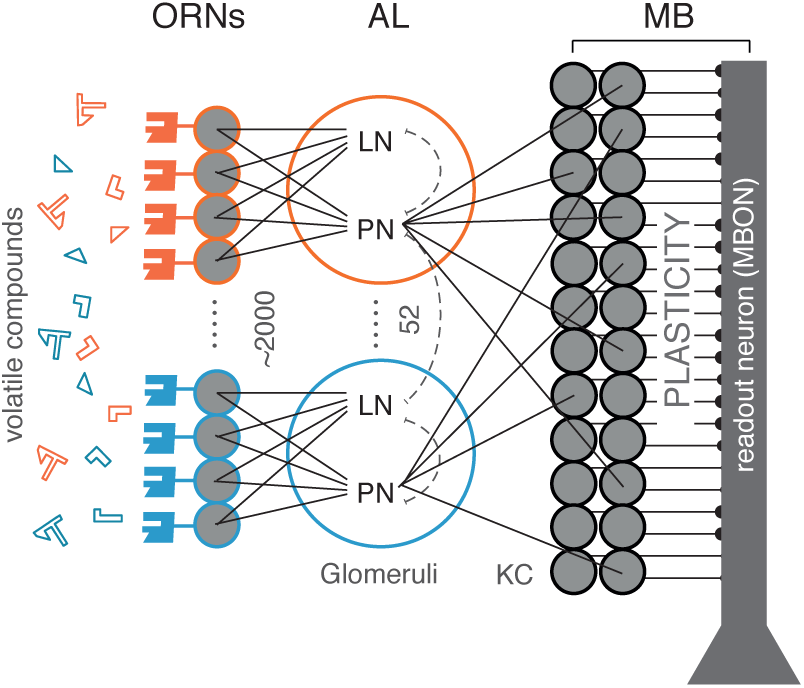
Spiking network model of the insect olfactory system. Olfactory receptor neurons (ORNs, *N* = 2080) at the antenna bind and respond to volatile odorant compounds. ORNs expressing the same (one of 52 different) genetic receptor type converge onto the same glumerus in the antennal lobe (AL). Each of the 52 glomeruli comprises one projection neuron (PN) and one local interneuron (LN). Each LN forms lateral inhibitory connections with all PNs. PNs randomly connect to a large population of Kenyon Cells (KC, *N* = 2000) where each KC receives input from on average ∼ 6 random PNs. All KCs project to a single MBON via plastic synapses.

### Sparse coding in space and time

The olfactory system transforms a dense olfactory code in the AL into a sparse stimulus code at the MB level. In the large population of KCs, a specific odor stimulus is represented by only a small fraction of all KCs (population sparseness) and each stimulus-activated KC responds with only a single or very few action potentials (temporal sparseness). In our model, temporal sparseness is achieved through the cellular mechanisms of spike-frequency adaptation (SFA) implemented at two levels of the system.

ORNs show clear stimulus response adaptation that has been attributed to the spike generating mechanism (27). Based on this experimental evidence we introduced a slow and weak SFA conductance in our model ORNs (see Materials and Methods). At the level of the MB, KCs have been shown to express strong SFA-mediating channels (18). This is matched by the SFA parameters of our model KCs (see Materials and Methods, (21)). As an effect of cellular adaptation in ORNs and KCs, odor stimulation (Fig 2 A) results in temporally precise and adaptive responses across all layers of the network (Fig 2 B). The effect of SFA implemented in ORNs is transitive and thus carries over to the postsynaptic PN and LN populations in agreement with experimental observations across species (28–31).

**Fig. 2.**
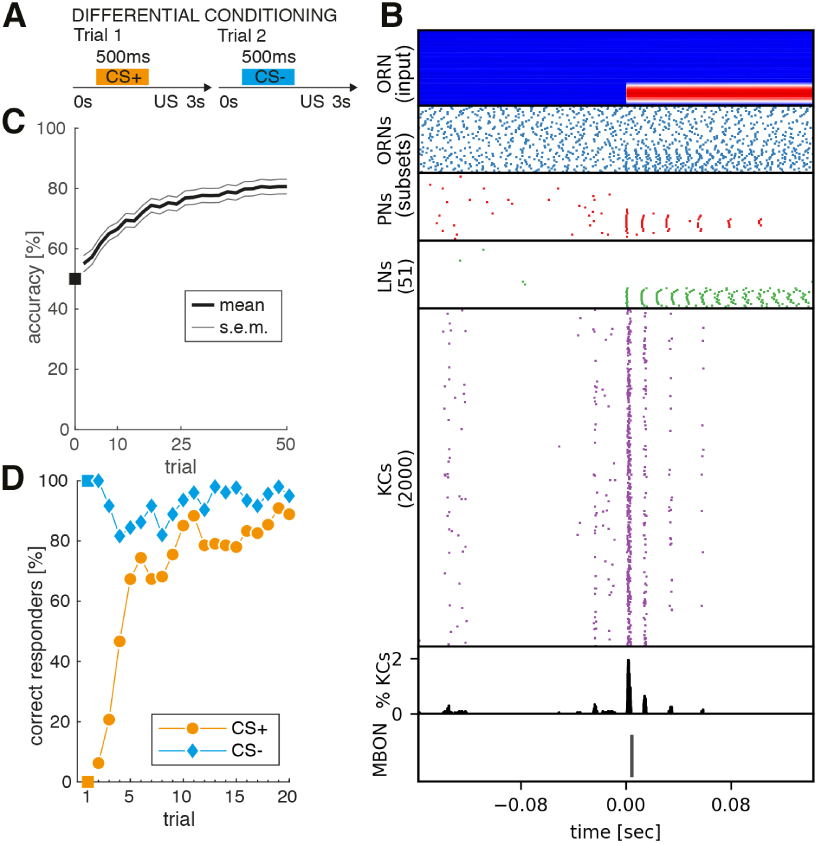
Rapid associative learning expressed in neuronal plasticity and conditioned response behavior. **A:** Sketch of differential conditioning protocol. In appetitive trials a first sensory cue (conditioned stimulus, CS+, orange) is paired with a reward (unconditioned stimulus, US). In aversive trials, a second sensory cue (CS-, blue) is paired with a punishment. Both trial types are presented randomized within blocks (see Materials and Methods). **B:** Sensory input and neuronal responses across all four circuit layers (ORN, AL, MB, MBON) in response to a CS+ odor presentation during the 10th training trial. Stimulus onset is at *t* = 0 s. From top to bottom: Model input is provided through independent noise current injection into the ORNs. The stimulus-induced input currents are clearly visible (hot colors) on top of the background noise for the subset of ORNs that are sensitive to the CS+ odor (stimulus profile). Stimulus response is clearly visible by an increase in the spiking activity across all neuron populations. For ORNs (blue) and PNs (red) relevant subsets of 60 and 35 neurons are shown. The population of 2000 KCs (magenta) show a temporal and spatial sparse odor response. Only 2% of all KCs are activated during a brief transient response following stimulus onset (black histogram). The MBON generates a single action potential in response to cue onset, which is the correct learned response to the CS+ odor. **C:** Learning performance of the MBON across *N* = 100 independent models as a function of the number of training trials. In any given trial the MBON response was correct if exactly one action potential was generated during CS+ presentation or if no action potential was generated during CS- presentation. **D:** The behavioral learning curve expresses the percentage of individuals that showed a correct behavior in the respective CS+ or CS- trial. The behavioral output is binary with either response or no response. The model triggers a response if the MBON generates one or more spikes.

In the KC population the background firing rate is very low (0.4 Hz). This is partially due to the outward SFA conductance and in agreement with experimental results (17). The KC population response is highly transitive where individual responding cells generate only a single or very few response spikes shortly after stimulus onset. This is in good qualitative and quantitative agreement with the temporal sparse KC spike responses measured in various species (17, 28, 32).

Population sparse stimulus encoding at the level of KCs is supported by two major factors. First, the sparse divergent-convergent connectivity between the PNs and the 20 times larger population of KCs is the anatomical basis for sparse odor representation (15, 20, 21, 33). Second, lateral inhibition mediated by the LNs in the AL (34) facilitates decorrelation of odor representations (34) and contributes to population sparseness (21). The sparse code in the KC population has been shown to reduce the overlap between different odor representations (35, 36) and consequently population sparseness is an important property of olfactory learning and plasticity models in insects (37–41). The system response to a single odor presentation in Fig. 2B) demonstrates the transformation of a dense olfactory code at the ORN and PN layers into a population sparse representation at the KC layer where less than < 2% of the total KC population is active at any time during stimulus presentation. This is in good agreement with quantitative estimates in the fruit fly (23, 42).

### Few-shot learning rapidly forms an associative memory of single cues with rewards

Many insects exhibit a rapid learning dynamics when trained in classical olfactory conditioning tasks. They typically acquire high retention scores (test accuracy > 60%) for a binary conditioned response (CR) behavior within only very few trials (e.g. (43–45)).

We here mimic a standard experimental lab protocol for differential conditioning (or acuity learning) to form associative memories and to generate a binary CR behavior by training our network (Fig. 1). Across successive learning trials we present two different odors in pseudo-random trial order (Fig. 2 A). Each trial constitutes a single odor presentation for 500 ms followed by a reinforcing stimulus (US) occurring shortly after the stimulus presentation. The CS+ odor is paired with a reward, the CS- odor with a punishment (see Materials and Methods). In order to establish a neural representation of the odor valence at the MB output (46–49) the MBON is trained (25, 26) to elicit exactly one action potential in response to the CS+ stimulus that is paired with the reward, and zero action potentials when the CS-stimulus is presented (see Materials and Methods). The system response to a single CS+ stimulus after nine conditioning trials is shown in Fig. 2 B.

In a first step we quantify the learning performance by considering the accuracy of the MBON response. MBON output is counted as correct if exactly one spike is generated during a CS+ trial and zero spikes during a CS- trial. The average accuracy over *N* = 100 independently trained model instances across successive trials is shown in Fig. 2 C. The learning dynamics shows a steep and steady increase indicating that an accurate memory is formed rapidly reaching up to 80% accuracy after 50 (25× CS+ and 25× CS-) training trials.

Next, we consider the behavioral learning curve, i.e. the acquisition of a binary CR behavior across successive learning trials. In each trial the model generates a behavioral response if the MBON produces one or more action potentials in response to the stimulus. No response is generated if the MBON remains silent. A CR is counted as correct if the MBON generates a response to the CS+ cue or no response to the CS- cue. The learning curve in Fig. 2 D represents the percentage of correctly responding individuals across *N* = 100 independently trained models. The untrained model, by default, does not generate any output spike, consequently 100% of the independent models correctly respond to CS- trials from the beginning (Fig. 2 D, blue). The red curve shows a rapid learning success where up to 70% of individuals generated the correct, appetitive CR to the CS+ stimuli within only 3 − 5 trials. The learning curve saturates after ∼ 10 trials with an asymptotic value of ∼ 80% correct responders. This reproduces the rapid learning dynamics of insects in classical conditioning experiments and fits qualitatively and quantitatively the CR behavior in honeybees (for review see (44)).

We conclude that our statically configured sensory network model with a single plastic readout neuron is capable to successfully form associative memories by few-shot learning, replicating the classical conditioning experiments in the typical lab situation. The computational mechanism of population sparseness implemented in our model increases discriminability of the two different stimuli supporting a rapid learning dynamics and a high accuracy of memory recall.

### Robust dynamic memory recall and odor-background segregation in complex sensory scenes

We now challenge our previously trained model (Fig. 2) in a novel task asking whether the already learned odor associations can be reactivated in a complex and dynamic olfactory scene. To this end we mimicked the encounter of odor filaments in a turbulent odor plume during a foraging flight (Fig. 3 A). For this we presented random sequences of non-overlapping olfactory cues within *T* = 10 s (see Materials and Methods). Each cue is of variable duration in the range between 100 ms and 500 ms. Odor identity of each cue is randomly assigned to either the CS+, CS- odor or one out of three additional background odors (Fig. 4 A). The use of non-overlapping cues follows the rationale that, in nature, filaments originating from different odors do not mix perfectly (50).

**Fig. 3.**
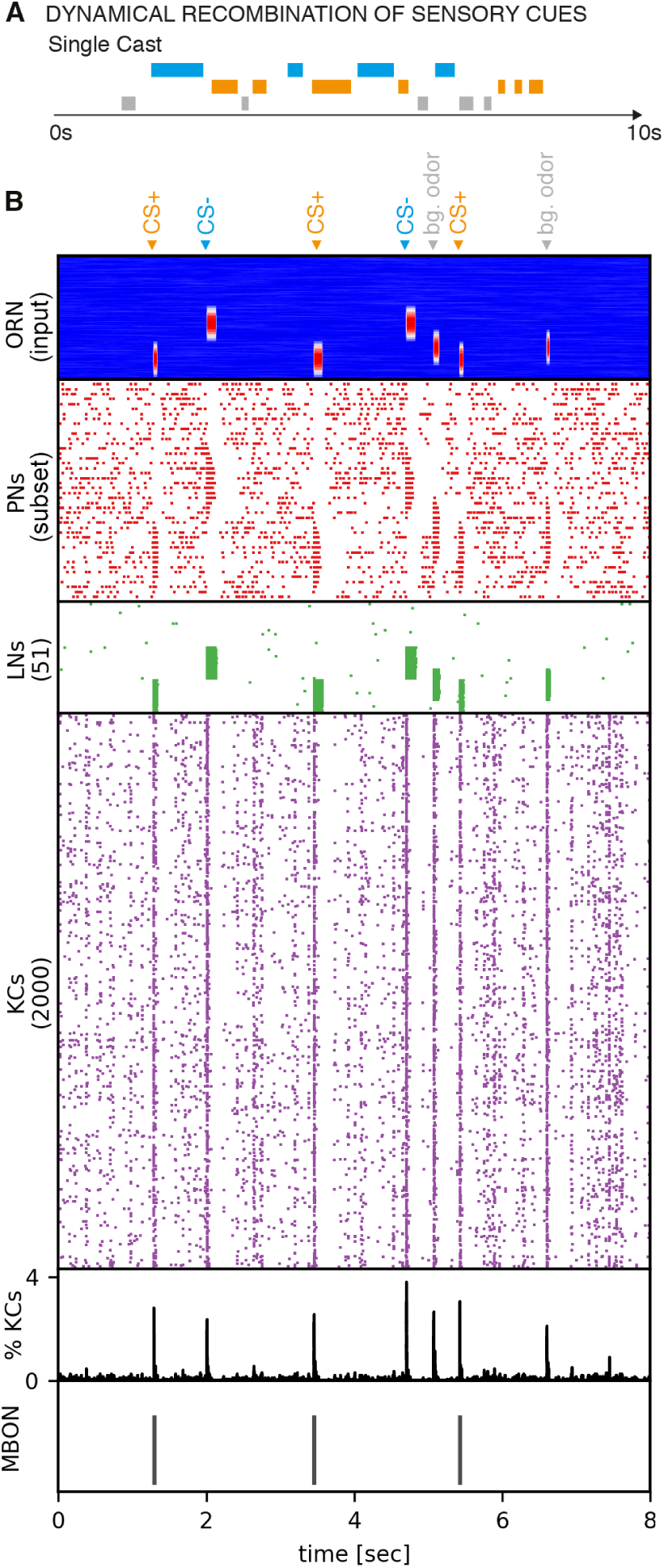
Recognition of valenced odor cues in complex dynamic scenes. **A:** Sketch of the dynamical memory recall task mimicking the sensory experience during a natural foraging flight. In each single trial of 10*s* duration the model encounters multiple (on average five) cues of different odor identities including the CS+ odor (orange), the CS- odor (blue) and background odors (gray). **B** Network response to one example input sequence made up of three CS+ cues, two CS- cues and two distractor cues (bg odor) as indicated at the top. The ORN input sequence indicates the fluctuating duration of odor cues. Transient PN and LN response porfiles faithfully represent individual odor cue onsets in time and odor identity across neuronal space. The KC population shows clear responses to all individual odor cues albeit with < 4% of activated cells at any time. The MBON correctly produced a single action potential in response to each of the three CS+ cues and zero output else.

**Fig. 4.**
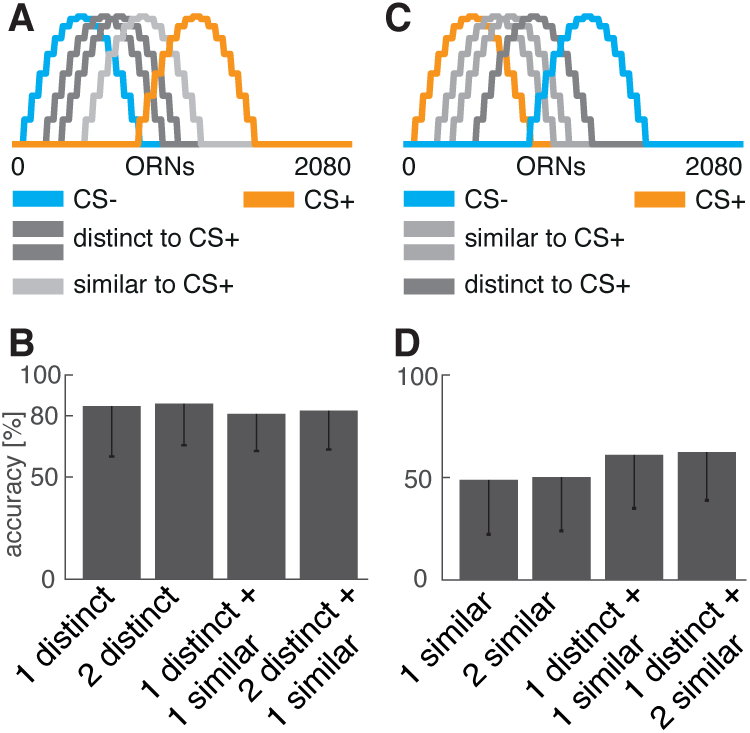
Performance in the dynamic memory recall task. **A:** Input activation profiles across ORNs for five different odors. The model had been previously trained in the differential conditioning protocol using the orange CS+ and the blue CS- odors. In the dynamic memory recall task (see text) background odors (gray) were presented as distractor cues along with CS+ and CS- cues. In the first task variant two background odor profiles were rather distinct from CS+ (dark gray) and one is more similar (light gray). **B:** Task accuracy across 200 trials in four different scenarios. A single trial consists of a random temporal sequence of sensory cue presentations during 10 s. Each single cue is of random duration between 100 − 500 ms. The single trial model response was correct if the MBON generated exactly as many action potentials as CS+ cues had been presented. The four task scenarios varied number (between one and three) and type (distinct vs. similar) of distractor cues as indicated. **C:** In the second task variant the same odors were used for cue presentation in a random temporal sequence. However, the model had been trained with reversed CS+/CS- odor contingency such that two distractor odors (light gray) are now more similar to the CS+ odor (orange). **D:** As in (B) but for reversed CS+/CS- odor contingency where distractor cues where overall more similar to the CS+ odor, increasing task difficulty.

The objective in this memory recall task is to correctly detect the occurrences of the positively valenced odor (CS+) by means of a single MBON action potential as model output while no output should be generated for all other cues (CS- or distractor odors). Fig. 3 B shows the system’s response to a singe random stimulus sequence where the MBON correctly generated a single action potential in response to each of three CS+ encounters. For quantification of task accuracy we considered the overall response to a given sequence to be correct if the number of action potentials generated by the readout neuron is equal to the number of CS+ cues.

For assessing model performance we systematically vary task difficulty by varying the number of possible background odors (between one and three) and their similarity with the CS+ odor (Fig. 4). In a first task variant background odor activation profiles are rather distinct from the CS+ odor and more similar to the CS- odor (Fig. 4 A). Accuracy of the model response is computed across 200 test sequences as shown in Fig. 4 B. We find that our previously trained model successfully generalizes to this new task with ∼ 80% accuracy for different sequence complexity in terms of identity and number of background odors. In a second task variant we reversed the odor contingency of the CS+ and CS- odors during initial differential conditioning. Thus, the reward predicting odor CS+ is now more similar to two of the background odors while similarity with the third background odor remains unchanged (Fig. 4 C). In this more challenging case accuracy reduces to ∼ 50 % of sequences for which the model produced the correct number of MBON output spikes. Note that the accuracy measure in Fig. 4 is based on the correct cumulative spike count during a complete trial of 10 s. The more similar a background odor stimulus profile is to the CS+ odor, the more likely the model will produce false positive (FP) action potentials in response to such a similar odor and thus a total spike count that is higher than the number of CS+ occurrences. This is reminiscent of the effect observed in insects and other animals in odor discrimination tasks where perceptually similar odors are more difficult to distinguish from previously learned CS+ odors than perceptually dis-similar odors during memory retention tests. This might be overcome if similar odors are used during the initial differential conditioning.

We conclude that our network model is able to recall previously learned neural representation of odors and signal their valence in a temporally dynamic setting where the rewarded and punished odors appear with up to five times shorter durations and within an unpredictable temporal cue sequence of previously unknown background odors. The models thus also solves the problem of odor vs. background segmentation under quasi natural conditions (51).

### Accumulation of sensory evidences informs motor control in foraging

We now consider the situation of foraging within a natural environment (Fig. 5 A). The objective is to locate the food source, which emits an attractive odor (CS+), by utilizing the sensory cues present in its turbulent odor plume. We show that cast & surge behavior can emerge by accumulation and exploitation of sensory evidence of sequentially experienced individual cues.

**Fig. 5.**
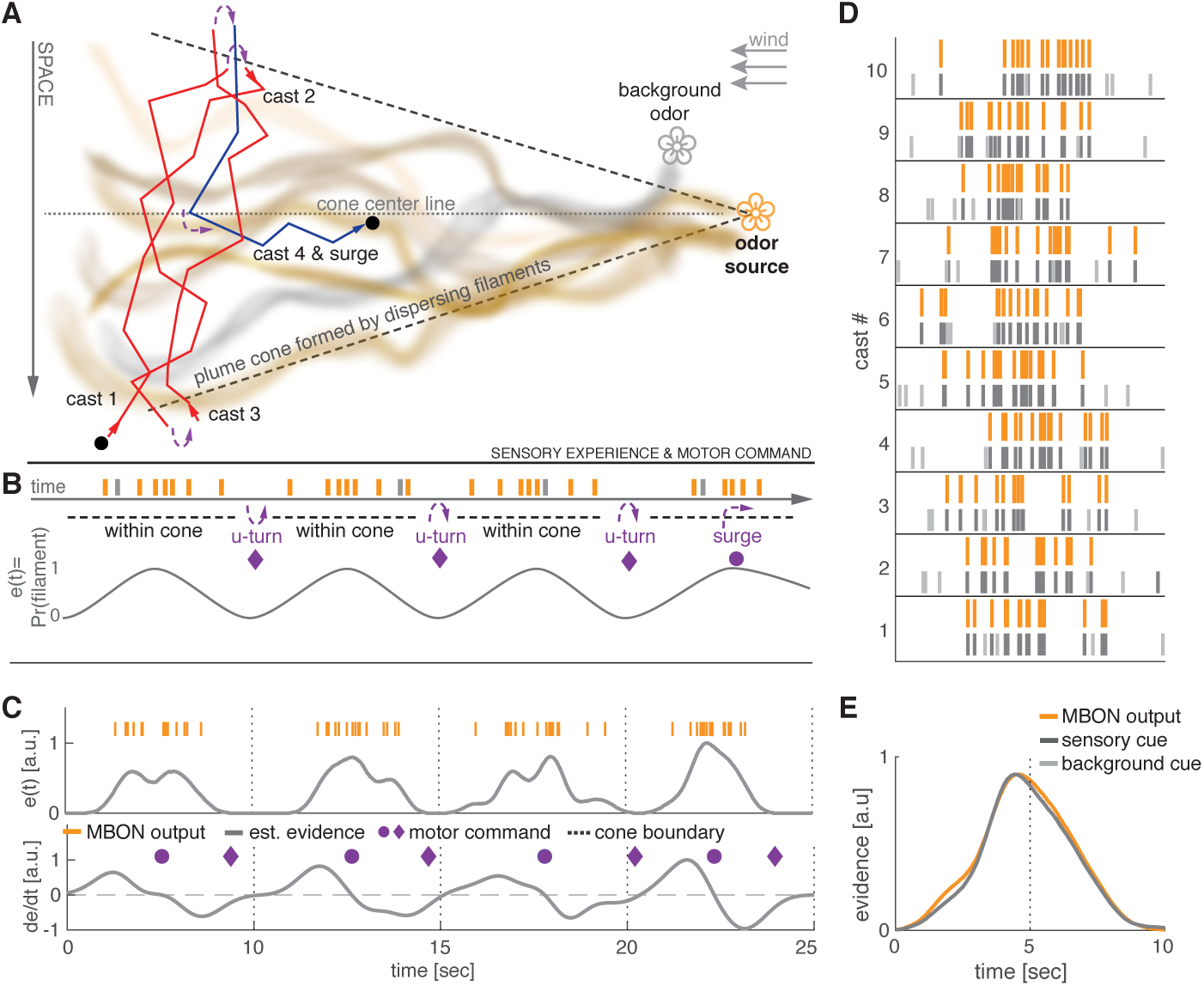
Dynamical sensory processing and motor control serving chemotaxis. **A:** Sketch of a typical olfactory experimental setup in a wind tunnel with a pleasant odor source (orange flower) and a second distractor source (gray flower). Due to turbulence the odor molecules emitted by a single source form dispersing, intermittent filaments within a cone-like boundary that constitutes the odor plume. The plume is modeled as Gaussian distributed filaments. A behaving model insect (here *Drosophila melanogaster*) performs stereotypic cast & surge behavior to locate the food source. This constitutes alternating between scanning crosswind and U-turning after running past the plume cone boundary where no filaments are present. Eventually, after several casts (here 3) it surges upwind until it loses track of the plume cone and starts over. **B:** Filament encounters during this behavior result in sequential brief on/off stimulations of the olfactory system. The probability of encountering filaments is *>* 0 within the plume and zero outside of the plume. Sensory evidence *e*(*t*) can be viewed as a likelihood function of filament encounters that increases towards the plume’s center line and is zero outside of the plume. The properties of this function can be used to generate optimal motor commands for chemotaxis. **C:** Evidence *e*(*t*) and derivative 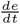 over 4 simulated successive casting trajectories estimated from the MBON spiking activity. *U-turn* motor commands (purple diamonds) are generated when *e*(*t*) runs below a fixed threshold (0.01) and *surge* motor commands (purple circles) are generated when 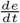 turns negative. The motor commands generated by the model match well with the theoretically optimal commands as sketched in panel B. **D:** Spiking activity of the MBON (orange) in response to 10 casting trajectories. The MBON reliably predicts the true sensory cues of positively valenced filaments (dark gray) and ignores background cues (light gray). **E:** Smooth PSTH computed over 10 casting trials recovers an accurate estimate of the true underlying sensory cue distribution simulated as Gaussian distribution.

For this task we assume that thin odor filaments within a cross-wind plane of the concentric odor plume are approximately Gaussian distributed. This is a reasonable assumption, particularly in a wind-tunnel setting with laminar flow as typically used in experimental settings (8, 52). When the insect performs a cast through the plume, it encounters filaments as short-lived discrete, sequential events where each encounter represents a single sensory cue (see sketch in Fig. 5B). Therefore, in our simulation of casting flights the agent encounters sequences of cues and distractors where cue onsets for the CS+ odor are drawn from a Gaussian distribution while distractor cue onsets appear uniformly distributed over time (see Materials and Methods). We further assume that the subject has already formed an association of food with the attractive odor, either through learning or through some genetically predetermined innate valence. To this end we again use the trained model from the classical conditioning task above (Fig. 2) without any further re-training.

We simulate 4 consecutive casting trajectories where the agent senses odor cues of sequentially experienced filament encounters. Ongoing accumulation of sensory evidence (Fig. 5C) by low-pass filtering of the readout neuron’s output assumes positive values shortly after entering the plume cone and further increases while approaching the plume’s center line. When travelling beyond the center line sensory evidence slowly decreases until the agent leaves the plume cone boundary. When sensory evidence drops to zero and after a fix delay, the agent initiates a U-turn motor command to perform another cross-wind cast.

Responses from our model’s readout neuron precisely follow the ground truth of CS+ odor cues as shown by 10 random casting trajectories in Fig. 5D. Performing analysis by averaging of sensory evidence across these 10 casting trajectories yields an average evidence (Fig. 5E) that faithfully resembles the underlying, true Gaussian profile of the simulated filaments.

We conclude that the model output provides an accurate and robust estimate of sensory evidence that can be used to reason about a plume’s spatial extend and center line. Both information are crucial to generate appropriate motor commands for U-turn and upwind surge behavior, necessary to successfully execute the cast & surge strategy. Apart from the existence of filaments inside a plume and absence outside a plume’s cone, our model does not make any specific assumption regarding the plume’s structure and statistics. It thus provides a generic mechanism implemented in a neural system to perform cast & surge behavior during foraging flights.

## Discussion

### Distinct functional roles for population and temporal sparse stimulus encoding

Population sparseness improves discriminability of different stimuli to facilitate associative learning. This has been demonstrated in theory and experiment (15, 20, 33, 36). We have shown, that our neural network model implements this feature in a biologically realistic way and our results confirm the functional role of population sparseness to support rapid and robust memory acquisition through associative learning. Experimental (36) and theoretical (20, 53) studies in the fruit fly strongly suggests that inhibitory feedback through the anterior paired lateral (APL) neuron improves population sparseness in the KC population. The APL is a GABAergic neuron that broadly innervates the KC population and likely receives input from KCs in the MB output region. Inhibitory feedback from MB output onto MB input has also been demonstrated in other species and blocking of feedback inhibition in the MB reduced population sparseness in the honeybee. Including an inhibitory feedback loop in our model would further improve robustness of population sparseness and thus not change our core findings.

Our model demonstrates how temporal sparseness can be exploited to generate short patterned signaling of cue identity. This enables perception of high temporal stimulus dynamics. In our model this is achieved independently of the duration of individual stimulus incidents and their distribution in time and makes temporally precise and robust sensory evidence available. It allows for the ongoing computation of derived estimates such as cue distributions or changes in cue density. Maintaining temporally sparse representations mechanistically supports the principle of compositionality (or Frege principle (22)), where an atomic stimulus entity is represented and can be learned by the readout neuron before processing this output. For example by estimation of densities or recombination with other entities to form composite perception or memory read-out. The temporal stimulus dynamics remains intact throughout the system even after learning of stimulus relevance. Thus valence is encoded with the same dynamics and faithfully captures occurrences of relevant cues. This allows compression of code to relevant stimuli while retaining full stimulus dynamics of the external world. Compression of code along the sensory processing pipeline is particularly relevant for small-brained animals like insects, which need to economize on their neuronal resources.

### Odor-background segregation: a joint effect of temporal and population sparse cue representation

The task presented in Fig. 3 implicitly addresses the issue of odor-background segregation. This refers to the problem that in nature cues of multiple odors of different sources are present, either in terms of mixtures or stimulus onset asynchrony due to turbulent conditions (50, 51). For behavior it is relevant to reliably isolate and detect the relevant cues from any background or distractor cues. The results presented in Fig. 4 show that this works nicely in our system. This is achieved by exploiting the joint effect of temporal and population sparseness. Optimal discrimination of cue representation is guarantied by population sparseness and temporal precision by means of temporal sparseness. Our plastic output neuron requires population sparseness for learning and the plasticity rule (25, 26) allows for temporally precise memory recall. We predict that our model can solve the challenge of odor-background segregation.

### Rapid learning within few trials

The ability of insects to quickly form associative memories after 3-5 trials has been demonstrated experimentally (44). However, in general few-shot learning remains a difficult task for computational models including insect inspired models (54). We find that, when compared with learning dynamics data of real insects (44) our model is able to show realistic learning dynamics that matches with the experimental observations. Due to frequent changes in the environment it might be a better strategy to trade-off fast and reasonable accurate learning against slow and high precision learning. Additionally, acquisition of training samples might be costly or they generally occur very sparsely.

Few-shot learning capabilities are also an active area of research in machine learning. Particularly current deep learning methods require massive amounts of training samples to successfully learn a classification model. For example, the popular benchmark data sets ImageNet and CIFAR10 for image classification contain 14 million and 60 million sample images, respectively. The *Google News dataset* used to train language models contains 100 billion words and learning to play the Space Invaders Atari game by deep reinforcement learning requires sampling of > 500.000 game frames from the environment. Clearly, these are numbers a biological organism cannot afford to accumulate. In fact few-shot learning likely is a fundamental skill for survival. We have demonstrated that our neurobiologically motivated approach using spike-based computations is capable to perform few-shot learning with similar speed as insects. We further showed that our model can transfer learned associations to novel, complex combinations that have not been part of the training data (transfer leaning).

### Innate vs. learned behavior

Cast & surge behavior belongs to the innate behavioral repertoire of air-borne insects and emerges from a set of sensori-motor reflexes (8). It can be considered as a base strategy which guarantees survival. The base system can be modulated and improved throughout an animal’s lifespan by experience-based learning. This is superior to alternative strategies that would solely rely on learning appropriate behaviors and thus require constant re-training as is the case in machine learning approaches. Here, we assumed that our readout neuron is tuned to a pleasant odor. In the present work this tuning is learned (adaptive process) in a classical conditioning task. However, a tuning can generally be learned by other mechanisms, e.g. reinforcement learning. We demonstrated that the existence of such a tuned neuron allows cast & surge foraging behavior to emerge.

There are other ways how such a tuned neuron can come about, for example due to genetically predetermined wiring or during development from larval to adult stage. The cast & surge behavior can be executed on innately valenced olfactory cues and our suggested model for motor control during cast & surge (Fig. 5A+B) also works for innate valenced stimuli. Learning is important to adapt behavior to changing environmental circumstances and associative learning provides a means to learn new valences on demand in such situations. Our model learns odor valences at the mushroom body output and it has been shown that MBONs signal odor valence (46– 49). We suggest that this valence is then used downstream to execute higher level functions of motor control. At this processing stage it might be integrated with innate valences and other necessary sensory modalities to form behavioral decisions.

### Implications for other sensory systems

Sparse stimulus encoding has been identified as a powerful principle used by higher order brain areas to encode and represent features of the sensory environment in invertebrate (17, 23, 42) and vertebrate (55–58) systems. Sensory systems with similar coding principles may share similar mechanisms when it comes to learning and multi-modal sensory integration. The mushroom body is a center for integration of multi-modal sensory information. Thus, our model can be extended to incorporate input from different sensory modalities. It is known that olfactory search and foraging strategies do not solely rely on olfactory cues, but require additional sensory information from at least visual cues and wind direction. Extending our model to include additional sensory processing systems for vision and wind direction can provide a comprehensive functional model to study foraging and navigation.

### Potential improvement through multiple readout neurons

Our current approach only comprises the simplest case of a single readout neuron. This model can be extended to multiple readout neurons. Different readout neurons can be tuned to different odors or groups of odorants. This would allow foraging for different types of food sources and further be useful for multi-modal sensory integration and learning of valences of multiple odors. Another way to use multiple readout neurons is to create an ensemble learning model. Particularly, one can perform bootstrap aggregation (bagging) to decrease variance of predictions. With this technique, multiple, independent readout neurons can be trained for the same target and their outputs are averaged to produce a single output. This approach can be useful when the level of noise increases due to different input models used to drive the network. Another possible extension is to use a single readout neuron to code for multiple odors by associating different number of action potentials to different odors (e.g. 2 or 3). The choice of model for the readout neuron and the plasticity rule allows to do this (25).

### Top-down motor control and lateral horn

The model currently lacks a neural implementation of sensory evidence integration and generation of motor commands. Integration of sensory evidence is modeled by low-pass filtering of the readout neuron’s spike train and its derivative is numerically estimated. In (59) it has been shown that a single compartment Hodgkin-Huxley neuron can operate in two computational regimes. One is more sensitive to input variance and acts like a differentiator while in the other regime it acts like an integrator. Similarly (60) has shown that the subthreshold current of neurons can encode the integral or derivative of their inputs based on their tuning properties. This and other suggested mechanistic implementations (e.g. (61)) could serve as basis for estimating the low-pass filtered sensory evidence and its derivative solely using neural computations. The initiation of a turning behavior based on a time-dependent evidence signal could be implemented e.g. through dis-inhibition of motor command neurons. The mechanism for U-turning could rely on either cell-intrinsic properties such as SFA where a neuron initiates a fast turning movement that decays with a fixed time constant, or through state-switching dynamics in neuronal populations.

### Relevance for machine learning and artificial intelligence

Learning and building artificial intelligent agents capable of interacting with their environment are major objectives in the field of machine learning (ML) and artificial intelligence (AI). Deep artificial neural networks (62) have demonstrated great success over the recent years. Particularly, in the domains of image recognition, natural language processing and deep reinforcement learning (63). Despite their success, when applied to agent-based systems, their major drawback becomes evident. They are very specific, single-purpose perceptual systems and poorly generalize to new tasks or changes in an agent’s environment (non-stationarities). A few methods to overcome this problem have been proposed, this includes retraining on new tasks, meta-learning and transfer-learning. In the context of deep learning this refers to the method of training a base network on features that are general to all tasks. Afterwards the pre-trained base network is used and the learned features are repurposed to only train a classification layer on the new tasks. However, it turned out that re-training brings up another weakness of deep neural networks, catastrophic forgetting (64). This term refers to the fact, that after a model has been trained on one task and gets re-trained on a second task, it will completely forget everything it has learned on the previous task. In this work we used a method similarly to the latter approach of transfer-learning but without any additional retraining and we used spike-based learning in an improved implementation (26) of the Multispike Tempotron (25). We predict that spike-based methods inspired by biological learning will become increasingly important for artificial intelligence.

## Materials and Methods

Code and data sets will be made available through our github profile at: https://github.org/nawrotlab

### Spiking network model

All neurons of the olfactory network are modeled as conductance-based leaky integrate-and-fire neurons with spike frequency adaptation (SFA). Specifically, the membrane potential follows the dynamical current balance equation 1. On threshold crossing a hard reset of the membrane potential is performed by 2. SFA is modeled as outward current by term 4 of equation 1. Strength of the adaptation current is modeled by a constant (*b*) decrease on each threshold crossing. Input to the model is modeled as direct, time-dependent current injection of shot-noise to all ORN neurons by the term *I*_*stim*_(*t*). All simulations of the network are carried out using BRIAN2 (65) simulator. The membrane potential of each neuron within a population is initialized randomly ∈ [*V*_*rest*_, *V*_*threshold*_]. To avoid any artifacts the network is brought to equilibrium by driving the network for 2 sec with background activity only before starting the actual simulation.

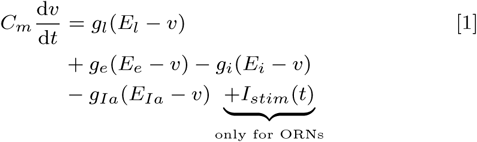

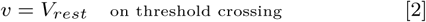

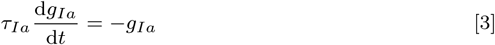

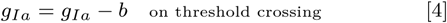

For this work the number of neurons within each layer and connectivity schemes are chosen to match the numbers found in the adult *Drosophila melanogaster* (14, 24). Our model comprises 2080 explicitly modeled olfactory recepter neurons (ORNs) organized in 52 different receptor types. ORNs of the same receptor type converge onto the same Glomerulus (52) by feedforward excitatory synapses. Each Glomerulus is formed by a projection neuron (PN) and local interneuron (LN). LNs provide lateral inhibition to all other PNs and LNs. PNs randomly project to a large population (2000) of Kenyon cells (KC) with excitatory synapses such that each KC on average receives input from 6 random PNs. This sparse random convergence implements population sparse responses. The single, plastic mushroom body output neuron is fully connected to all KCs.

We used the cellular mechanism of spike-frequency adaptation (SFA) to achieve temporal sparseness. ORNs are configured to have slow and weak spike-frequency adaptation in accordance with experimental findings (27, 30). For PNs and LNs SFA has been turned off and KCs are set to produce fast and strong adaptation currents (18, 66). The property of temporal sparseness can also be achieved by an alternative implementation through feedback inhibition as proposed by (53) and (67).

The synaptic weights of all connections within the network have been manually determined such that an average background firing rate of 8 − 10 Hz is achieved in the LN population.

### Stimulus response profile of ORNs

The stimulus response profile of ORNs is determined by the ORN tuning curves. We follow a similar method as used in (21) where cyclical tuning over receptor types is modeled as half period sine waveforms. Our model comprises *N*_*type*_ = 52 receptor types and supports 52 different stimuli (e.g. different odors). Where *k*_*type*_ refers to the receptor type index (∈[0, 51]) and *k*_*odor*_ to the stimulus index (∈[0, 51]). *N*_*orn*_ = 15 determines the number of receptor types activated by a stimulus. The tuning strength *r* of the ORNs can be computed as 0.5 cycle of a sine wave with peak amplitude *r*_*max*_ = 1. In the present work all tuning profiles are normalized to have a peak amplitude of 1.

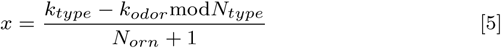

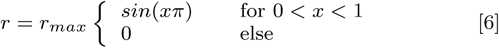

### Model input

Input to the mushroom body model is modeled as time-dependent, direct current injection into all ORN neurons. In the absence of any stimuli ORNs exhibit spontaneous activity (27). The model input thus consists of spontaneous background activity and stimulus related activity. To generate the background activity, a current time-series is generated for each ORN by simulating shot noise. For each ORN neuron, background activity events are generated from a Poisson process with high rate (*λ* = 300) (independent Poisson processes are drawn for each individual neuron). Events of the Poisson process are filtered by a low-pass filter with *τ* = 0.6 sec. Using this shot-noise model is consistent with experimental findings of odor transduction at the ORNs (27). To induce stimulus related activity to this time-series of ORN *j* it is multiplied point-wise with a stimulaton protocol time-series *s*_*j*_ (*t*) which is rescaled by a constant determined by the tuning strength (*r*_*j*_ ∈ [0, 1]) to the specific odor of the ORN. This results in a current time series where during stimulus the current magnitude is increased proportional to the ORNs tuning strength and otherwise remains at the magnitude of the background activity.

We define a stimulation protocol function *s*(*t*), which is a step function taking on the value 1 at all time points *t* where a stimulus or sensory cue is active. For each ORN a rescaled instance of the stimulation protocol is defined as *s*_*j*_ (*t*) = *r*_*j*_ *s*(*t*), where the scaling parameter *r*_*j*_ ∈ [0, 1] is given by the stimulus response profile (eq. 6) of the ORN to the specific stimulus.

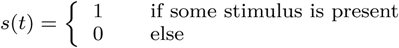

### Sequences of sensory cues

Each sequence has a duration of 10 seconds. Sequences of sensory cues are generated by drawing the total number of cues within a single sequence from a Poisson distribution with mean *λ* = 8. Onset times of the cues between 0 and 10 seconds are drawn from a random uniform distribution and it is assured that there is no temporal overlapp between cues. A stimulus relates to a single sensory cue and its duration is drawn uniformly between [1, 200] milliseconds. Finally, each sensory cue is associated with a random odor drawn from a fixed set of possible odors (random sampling with replacement and equal probability). This results in sequences with random number of sensory cues, random onset, random duration and randomized odor and distractor combinations.

### Model of sensory cues within (gaussian) plume

The same procedure is used as above to simulate the experience of sensory cues during a single casting trajectory within a turbulent odor plume. The number of pleasant cues experienced in a casting trajectory is drawn from a Poisson distribution with mean *λ* = 14. The cue onset times are drawn from a gaussian distribution with *µ* = 5, *σ* = 1.5. The number of distractor cues is drawn from a Poisson distribution with mean *λ* = 5 and are distributed uniformly in time. Duration of both, pleasant and distractor cues, is drawn uniformly between [100, 500] ms. In total 200 different casting trajectories have been generated using this procedure.

### Readout Neuron & Learning rule

To fit the readout neuron to the stimuli such that it generates 1 spike for pleasant odor stimuli (CS+) and 0 spikes for any other stimuli (CS-) we use a modified implementation of the Multispike Tempotron (25, 26). Thus, the readout neuron is modeled as voltage-based leaky integrate-and-fire neuron with soft reset following the dynamical equation 7. Incoming spikes evoke exponentially decaying post-synaptic potentials. When the membrane potential reaches the spiking threshold at some time *t*_0_ an output spike is generated and the membrane potential is reset by the last term of equation 7.

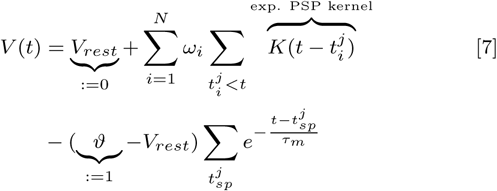

The dynamical equation can be decomposed into two parts, the unreset sub-threshold potential *V*_0_(*t*) (eq. 8) minus the remaining terms for the soft-reset. The neuron is trained to generate 1 spike for pleasant odor stimuli (CS+1) and 0 spikes for any other stimuli (CS-). To fit the desired neural code, a training step is performed after each stimulus presentation. A training step is performed only if the number of spikes generated in response to a stimulus was not correct. The training target is given by the difference between number of output spikes the model generated and the number of output spikes associated with the stimulus. We denote the desired critical threshold value, the voltage value that generates *d* = 1 spike, as *ϑ*^∗^ and the time point where this voltage value is reached by *t*^∗^ (more generally: the critical threshold value to generate *d* spikes). We briefly sketch the idea and intuition of the Multispike Tempotron learning rule. For detailed derivation of the rule we refer to (25) and the section *The ϑ*^∗^ *gradient*. The Multispike Tempotron training algorithm works by differentiating the membrane potential of the the critical threshold wrt. to the synaptic weights 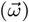. This can be done since *ϑ*^∗^ is a regular voltage value, that can be expressed by the neuron’s dynamical equation Eq. (7), with the special identities shown in equation 10. This allows to take the full derivative as shown in equation 11.

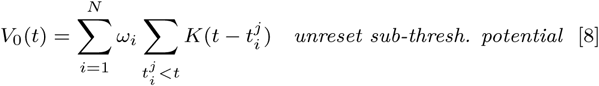

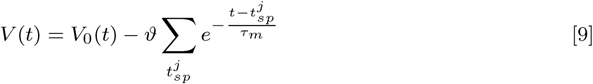

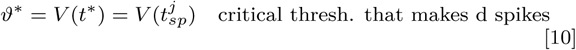

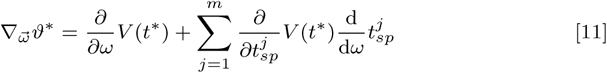

The gradient of the critical threshold with respect to a single synapse *i* is given by equation 12.

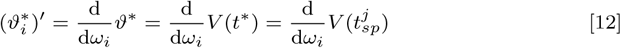

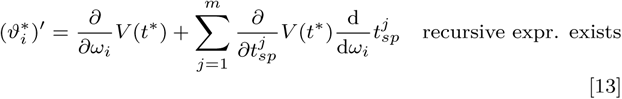

## ACKNOWLEDGMENTS

This research is supported by the German Research Foundation (grant no. 403329959 to MN) within the Research Unit ‘Structure, Plasticity and Behavioral Function of the Drosophila Mushroom Body’ (DFG-FOR 2705, www.uni-goettingen.de/en/601524.html).

